## Supplemental materials for "Recurrent Mutations Drive the Rapid Evolution of Pesticide Resistance in the Two-spotted Spider Mite *Tetranychus urticae*"

Table S1 Population information of the two-spotted spider mite *Tetranychus* *urticae* used in this study

Table S2 Mutant allele frequency (Sanger / pooled sequencing) and susceptibility of the two-spotted spider mites *Tetranychus* *urticae* to cyetpyrafen

Table S3 Laboratory selection and cross-resistance study of *Tetranychus* *urticae*

Table S4 Sequencing information and genetic diversity of 22 populations of the two-spotted spider mite *Tetranychus* *urticae*

Table S5 Distribution of haplotypes carrying amino acid mutations in populations

Table S6 Correlation between eight mutations on SDH genes and level of resistance (survival percentages under 1000 mg/L)

Fig. S1 Line chart of nucleotide diversity (a - b) and Tajima’D values (c - d) along chromosomes near the target gene of cyetpyrafen

Fig. S2 Scatter plot of average genome-wide nucleotide diversity (π) and Tajima's D per population

Fig. S3 Haplotypes of sdhB (a) and sdhD (b) genes

Fig. S4 Correlation between allele frequencies genotyped by Sanger sequencing and pool-seq.

**Table S1 Population information of the two-spotted spider mite *Tetranychus* *urticae* used in this study**

| **Population** | **Date** | **Location** | **Host** | **Pool-seq** | **Sanger-seq** | **Bioassay** | **Data source** |
| --- | --- | --- | --- | --- | --- | --- | --- |
| BJCP1 | 2013.04.25 | Changping District, Beijing | strawberry |  | Y |  | This study |
| BJPG1 | 2013.05.23 | Pinggu District, Beijing | strawberry |  | Y |  | This study |
| BJHD1 | 2017.03.13 | Haidian District, Beijing | eggplant |  | Y |  | This study |
| SXYQ | 2017.03.14 | Yangquan City, Shanxi Province | strawberry | Y |  |  | This study |
| AHHN | 2017.03.16 | Huainan City, Anhui Province | strawberry | Y |  |  | This study |
| SDRZ | 2017.03.22 | Rizhao City, Shandong Province | strawberry | Y | Y |  | This study |
| SDSG1 | 2017.03.29 | Shouguang City, Shandong Province | pepper |  | Y |  | This study |
| HNHK | 2017.03.30 | Haikou City, Hainan Province | kidney beans | Y | Y |  | This study |
| SCCD1 | 2017.03.30 | Chengdu City, Sichuan Province | strawberry | Y | Y |  | This study |
| SHPD | 2017.04.13 | Pudong District, Shanghai | strawberry | Y | Y |  | This study |
| HNCS1 | 2017.04.02 | Changsha City, Hunan Province | strawberry | Y | Y |  | This study |
| BJTZ1 | 2017.04.24 | Tongzhou District, Beijing | strawberry |  | Y |  | This study |
| BJPG2 | 2017.05.05 | Pinggu District, Beijing | strawberry |  | Y |  | This study |
| JXNC | 2017.05.08 | Nanchang City, Jiangxi Province | strawberry | Y |  |  | This study |
| BJCP2 | 2017.07.26 | Changping District, Beijing | strawberry |  | Y |  | This study |
| BJDX | 2018 | Daxing District, Beijing | strawberry |  |  | Y | Gong et al. 2018 |
| BJDX1 | 2018.02 | Daxing District, Beijing | strawberry |  |  | Y | Chen et al. 2019 |
| BJDX2 | 2018.04 | Daxing District, Beijing | strawberry |  |  | Y | Chen et al. 2019 |
| BJFS | 2018.04 | Fangshan District, Beijing | strawberry |  |  | Y | Chen et al. 2019 |
| ZJXS1 | 2018.04 | Hangzhou City, Zhejiang Province | strawberry |  | Y | Y | Chen et al. 2019 |
| BJSY | 2018.05 | Shunyi District, Beijing | strawberry |  |  | Y | Chen et al. 2019 |
| ZJWX | 2018.05 | Wuxing City, Zhejiang province | strawberry |  |  | Y | Chen et al. 2019 |
| ZJHZ | 2020.04.14 | Hangzhou City, Zhejiang Province | strawberry |  |  | Y | This study |
| HBBD | 2020.04.15 | Baoding City, Hebei Province | strawberry |  |  | Y | This study |
| HNZZ | 2020.04.15 | Zhengzhou City, Henan Province | strawberry |  | Y | Y | This study |
| SDSG2 | 2020.04.16 | Shouguang City, Shandong Province | kidney beans |  | Y | Y | This study |
| BJDX3 | 2020.04.17 | Daxing District, Beijing | strawberry |  |  | Y | This study |
| BJYQ | 2020.04.17 | Yanqing District, Beijing | strawberry |  |  | Y | This study |
| SXAK | 2020.04.17 | Ankang City, Shaanxi Province | strawberry |  | Y | Y | This study |
| ZJJX | 2020.04.17 | Jiaxing City, Zhejiang Province | strawberry |  |  | Y | This study |
| ZJNB | 2020.04.17 | Ningbo City, Zhejiang Province | strawberry |  |  | Y | This study |
| NMHH1 | 2020.04.18 | Hohhot City, Inner Mongolia Autonomous Region | strawberry |  |  | Y | This study |
| SDQD | 2020.04.20 | Qingdao City, Shandong Province | strawberry |  | Y | Y | This study |
| YNKM1 | 2020.04.23 | Kunming City, Yunnan Province | rose |  | Y | Y | This study |
| BJHD2 | 2020.04.26 | Haidian District, Beijing | strawberry |  | Y | Y | This study |
| LNSY | 2020.05.22 | Shenyang City, Liaoning Province | strawberry |  | Y | Y | This study |
| YNKM2 | 2021.11.03 | Kunming City, Yunnan Province | rose |  | Y | Y | This study |
| BJCP4 | 2021.04.14 | Changping District, Beijing | strawberry | Y | Y | Y | This study |
| SDSG3 | 2021.04.18 | Shouguang City, Shandong Province | cucumber | Y | Y | Y | This study |
| SDSG4 | 2021.04.18 | Shouguang City, Shandong Province | eggplant |  | Y | Y | This study |
| BJCP5 | 2021.04.27 | Changping District, Beijing | strawberry |  |  | Y | This study |
| BJPG3 | 2021.04.27 | Pinggu District, Beijing | strawberry |  | Y |  | This study |
| SCCD2 | 2021.06.12 | Chengdu City, Sichuan Province | eggplant |  | Y |  | This study |
| NMHH2 | 2021.06.13 | Hohhot City, Inner Mongolia Autonomous Region | kidney beans | Y | Y | Y | This study |
| SDQZ | 2021.06.13 | Qingzhou City, Shandong Province | eggplant | Y | Y | Y | This study |
| NXGY2 | 2021.06.27 | Guyuan City, Ningxia Province | cucumber |  | Y | Y | This study |
| BJHD3 | 2021.07.23 | Haidian District, Beijing | corn |  | Y |  | This study |
| SDSG5 | 2021.07.07 | Shouguang City, Shandong Province | eggplant |  | Y | Y | This study |
| SDSG6 | 2021.07.07 | Shouguang City, Shandong Province | pepper |  | Y | Y | This study |
| BJTZ2 | 2021.04.30 | Tongzhou District, Beijing | strawberry |  | Y |  | This study |
| BJMY | 2024.01.30 | Miyun District, Beijing | strawberry |  |  | Y | This study |
| SDSG7 | 2024.01.30 | Shouguang City, Shandong Province | cucumber |  |  | Y | This study |
| BJHD4 | 2024.02.24 | Haidian District, Beijing | strawberry |  | Y | Y | This study |
| BJDX5 | 2024.03.15 | Daxing District, Beijing | strawberry |  | Y | Y | This study |
| BJDX6 | 2024.03.15 | Daxing District, Beijing | strawberry | Y | Y | Y | This study |
| BJDX7 | 2024.01.30 | Daxing District, Beijing | strawberry |  | Y | Y | This study |
| GXNN | 2024.03.31 | Naning City, Guangxi Province | strawberry | Y | Y | Y | This study |
| GZGY | 2024.04.03 | Guiyang City, Guizhou Province | strawberry |  | Y |  | This study |
| HNCS2 | 2024.01.29 | Changsha City, Hunan Province | strawberry | Y | Y | Y | This study |
| JXYC | 2024.03.10 | Yichun City, Jiangxi Province | strawberry |  | Y |  | This study |
| LNDD | 2024.03.10 | Dandong City, Liaoning Province | strawberry | Y | Y | Y | This study |
| QHHD | 2024.02.29 | Haidong City, Qinghai Province | strawberry | Y | Y | Y | This study |
| SCCD3 | 2024.04.08 | Chengdu City, Sichuan Province | strawberry |  | Y |  | This study |
| SDSG8 | 2024.03.12 | Shouguang City, Shandong Province | strawberry |  | Y |  | This study |
| SDWF | 2024.04.10 | Weifang City, Shandong Province | eggplant |  | Y |  | This study |
| YNKM3 | 2024.04.07 | Kunming City, Yunnan Province | strawberry |  | Y |  | This study |
| YNKM4 | 2024.04.04 | Kunming City, Yunnan Province | pepper |  | Y |  | This study |
| YNYX | 2024.03.5 | Yuxi City, Yunnan Province | strawberry | Y | Y | Y | This study |
| ZJHZ2 | 2024.03.15 | Hangzhou City, Zhejiang Province | strawberry | Y | Y | Y | This study |
| ZJHZ3 | 2024.03.15 | Hangzhou City, Zhejiang Province | strawberry | Y | Y | Y | This study |
| LabS | 2018.04 | Hangzhou City, Zhejiang Province | strawberry | Y |  | Y | This study |
| LabR | 2018.04 | Hangzhou City, Zhejiang Province | strawberry | Y | Y | Y | This study |

Y.-J. Gong et al., Toxicity and field control efficacy of the new acaricide SYP-9625 to the two-spotted spider mite (Tetranychus urticae koch). Agrochemicals 56, 561-563 (2017).

J. C. Chen et al., Field-evolved resistance and cross-resistance of the two-spotted spider mite, Tetranychus urticae, to bifenazate, cyenopyrafen and SYP-9625. Experimental and Applied Acarology (2.38) 77, 545-554 (2019).

**Table S2 Mutant allele frequency (Sanger / pooled sequencing) and susceptibility of the two-spotted spider mites *Tetranychus* *urticae* to cyetpyrafen.** #, from Gong et al. 2018; $, from Chen et al., 2019. Sample size (sdhB/sdhD). Frequency of mutant individuals (carrying any of the mutation) is calculated from Sanger sequencing data of all mutations except for H146Q, S212I, A285S, D116E, and R119P mutations. SR, survival rate (%) under 1000 mg/L. LC_50_ (mg/L), number of proportions shows mortality treated by 32000 mg/L.

| **Population** | **Year** | ***B*_H146Q** | ***B_*S212I** | ***B_H258Y*** | ***B_*I260T** | ***B_*I260V** | ***B_*A285S** | ***D_*D116G** | ***D_*D116E** | ***D_*D116N** | ***D_*R119C** | ***D_*R119G** | ***D_*R119L** | ***D_*R119P** | ***D_*R119H** | ***D_P120L*** | **No. mutation** | **Mutant indiv. %** | **LC_50_ (mg/L)** | **SR** |
| --- | --- | --- | --- | --- | --- | --- | --- | --- | --- | --- | --- | --- | --- | --- | --- | --- | --- | --- | --- | --- |
| BJCP1 | 2013 | - | - | 0/- | 0/- | 0/- | 0/- | 0/- | 0/- | 0/- | 0/- | 0/- | 0/- | 0/- | 0/- | 0/- | 0 | 0 | - | - |
| BJPG1 | 2013 | - | - | 0/- | 0/- | 0/- | 0/- | 0/- | 0/- | 0/- | 0/- | 0/- | 0/- | 0/- | 0/- | 0/- | 0 | 0 | - | - |
| BJCP2 | 2017 | - | - | 0/- | 0/- | 0/- | 0/- | 0/- | 0/- | 0/- | 0/- | 0/- | 0/- | 0/- | 0/- | 0/- | 0 | 0 | - | - |
| BJHD1 | 2017 | - | - | 0/- | 0/- | 0/- | 0/- | 0/- | 0/- | 0/- | 0/- | 0/- | 0/- | 0/- | 0/- | 0/- | 0 | 0 | - | - |
| BJPG2 | 2017 | - | - | 0/- | 0/- | 0/- | 0/- | 0/- | 0/- | 0/- | 0/- | 0/- | 0/- | 0/- | 0/- | 0/- | 0 | 0 | - | - |
| SXYQ | 2017 | -/0 | -/0 | -/0 | -/0 | -/0 | -/0 | -/0 | -/0 | -/0 | -/0 | -/0 | -/0 | -/0 | -/0 | -/0 | 0 | - | - | - |
| AHHN | 2017 | -/0 | -/0 | -/0 | -/0 | -/0 | -/0 | -/0 | -/0 | -/0 | -/0 | -/0 | -/0 | -/0 | -/0 | -/0 | 0 | - | - | - |
| JXNC | 2017 | -/0 | -/0 | -/0 | -/0 | -/0 | -/0 | -/0 | -/0 | -/0 | -/0 | -/0 | -/0 | -/0 | -/0 | -/0 | 0 | - | - | - |
| BJTZ1 | 2017 | - | - | 0/- | 0/- | 0/- | 0/- | 0/- | 0/- | 0/- | 0/- | 0/- | 0/- | 0/- | 0/- | 0/- | 0 | 0 | - | - |
| HNCS1 | 2017 | -/0 | -/0 | 0/0 | 0/0 | 0/0 | 0/0 | 0/0 | 0/0 | 0/0 | 0/0 | 0/0 | 0/0 | 0/0 | 0/0 | 0/0 | 0 | 0 | - | - |
| HNHK | 2017 | -/0 | -/0 | 0/0 | 0/0 | 0/0 | 0/0 | 0/0 | 0/0 | 0/0 | 0/0 | 0/0 | 0/0 | 0/0 | 0/0 | 0/0 | 0 | 0 | - | - |
| SCCD1 | 2017 | -/0 | -/0 | 0/0 | 0/0 | 0/0 | 0/0 | 0/0 | 0/0 | 0/0 | 0/0 | 0/0 | 0/0 | 0/0 | 0/0 | 0/0 | 0 | 0 | - | - |
| SDRZ | 2017 | -/0 | -/0 | 0/0 | 0/0 | 0/0 | 1.9/33 | 0/0 | 0/0 | 0/0 | 0/0 | 0/0 | 0/0 | 0/0 | 0/0 | 0/0 | 1 | 0 | - | - |
| SDSG1 | 2017 | - | - | 0/- | 0/- | 0/- | 0/- | 0/- | 0/- | 0/- | 0/- | 0/- | 0/- | 0/- | 0/- | 0/- | 0 | 0 | - | - |
| SHPD | 2017 | -/0 | -/0 | 0/0 | 0/0 | 0/0 | 0/0 | 0/0 | 0/0 | 0/0 | 0/0 | 0/0 | 0/0 | 0/0 | 0/0 | 0/0 | 0 | 0 | - | - |
| BJDX# | 2018 | - | - | - | - | - | - | - | - | - | - | - | - | - | - | - | - | - | 9.64 | - |
| BJDX1$ | 2018 | - | - | - | - | - | - | - | - | - | - | - | - | - | - | - | - | - | 2.51 | - |
| BJDX2$ | 2018 | - | - | - | - | - | - | - | - | - | - | - | - | - | - | - | - | - | 2.15 | - |
| BJFS$ | 2018 | - | - | - | - | - | - | - | - | - | - | - | - | - | - | - | - | - | 4.15 | - |
| ZJXS1$ | 2018 | - | - | 0/- | 0/- | 0/- | 0/- | 0/- | 0/- | 0/- | 0/- | 0/- | 0/- | 0/- | 0/- | 0/- | 0 | 0 | 21.55 | 0 |
| BJSY$ | 2018 | - | - | - | - | - | - | - | - | - | - | - | - | - | - | - | - | - | 1.4 | - |
| ZJWX$ | 2018 | - | - | - | - | - | - | - | - | - | - | - | - | - | - | - | - | - | 6.54 | - |
| HNZZ | 2020 | - | - | 0/- | 0/- | 0/- | 0/- | 0/- | 0/- | 0/- | 0/- | 0/- | 0/- | 0/- | 1.79/- | 0/- | 1 | 0 | 1.31 | 0 |
| BJDX3 | 2020 | - | - | - | - | - | - | - | - | - | - | - | - | - | - | - | - | - | 1.89 | - |
| BJHD2 | 2020 | - | - | 0/- | 0/- | 0/- | 0/- | 0/- | 0/- | 0/- | 0/- | 0/- | 0/- | 85.71/- | 0/- | 0/- | 1 | 0 | 12.95 | 0 |
| BJYQ | 2020 | - | - | - | - | - | - | - | - | - | - | - | - | - | - | - | - | - | 1.78 | - |
| HBBD | 2020 | - | - | - | - | - | - | - | - | - | - | - | - | - | - | - | - | - | 2.8 | - |
| LNSY | 2020 | - | - | 0/- | 0/- | 0/- | 0/- | 0/- | 0/- | 0/- | 25.0/- | 0/- | 3.57/- | 0/- | 0/- | 0/- | 2 | 27.27 | 4.04 | 0 |
| NMHH1 | 2020 | - | - | - | - | - | - | - | - | - | - | - | - | - | - | - | - | - | 1.72 | - |
| SDQD | 2020 | - | - | 0/- | 0/- | 0/- | 0/- | 0/- | 0/- | 3.45/- | 0/- | 0/- | 0/- | 0/- | 0/- | 0/- | 0 | 3.45 | 2.11 | 0 |
| SDSG2 | 2020 | - | - | 0/- | 12.5/- | 0/- | 0/- | 66.7/- | 0/- | 0/- | 0/- | 0/- | 0/- | 0/- | 0/- | 0/- | 2 | 100 | 14.7% | 85.7 |
| SXAK | 2020 | - | - | 0/- | 0/- | 0/- | 0/- | 0/- | 0/- | 0/- | 0/- | 0/- | 0/- | 0/- | 0/- | 0/- | 0 | 0 | 2.29 | 0 |
| YNKM1 | 2020 | - | - | 0/- | 3.8/- | 0/- | 0/- | 0/- | 0/- | 0/- | 0/- | 0/- | 0/- | 0/- | 0/- | 0/- | 1 | 5.56 | 7.57 | 0 |
| ZJHZ | 2020 | - | - | - | - | - | - | - | - | - | - | - | - | - | - | - | - | - | 4.27 | - |
| ZJJX | 2020 | - | - | - | - | - | - | - | - | - | - | - | - | - | - | - | - | - | 1.72 | - |
| ZJNB | 2020 | - | - | - | - | - | - | - | - | - | - | - | - | - | - | - | - | - | 1.97 | - |
| BJCP4 | 2021 | -/2.13 | -/0 | 0/0 | 1.9/0 | 44.2/54.5 | 0/0 | 0/0 | 0/0 | 0/0 | 78.1/82.8 | 0/0 | 0/0 | 0/0 | 0/0 | 0/0 | 4 | 100 | 13546.15 | 74.6 |
| BJHD3 | 2021 | - | - | 0/- | 0/- | 27.6/- | 0/- | 0/- | 0/- | 0/- | 0/- | 0/- | 0/- | 65.0/- | 0/- | 0/- | 2 | 93.10 | - | - |
| BJCP5 | 2021 | - | - | - | - | - | - | - | - | - | - | - | - | - | - | - | - | - | > 13000 | - |
| BJPG3 | 2021 | - | - | 0/- | 0/- | 100/- | 0/- | 0/- | 0/- | 0/- | 4.0/- | 0/- | 14.0/- | 0/- | 0/- | 0/- | 2 | 100 | - | - |
| BJTZ2 | 2021 | - | - | 0/- | 0/- | 80.0/- | 0/- | 0/- | 0/- | 0/- | 12.5/- | 0/- | 0/- | 0/- | 0/- | 0/- | 2 | 87.5 | - | - |
| NMHH2 | 2021 | -/0 | -/0 | 0/0 | 0/0 | 0/5.5 | 0/0 | 0/0 | 0/0 | 0/0 | 0/0 | 0/0 | 0/0 | 0/0 | 0/0 | 0/0 | 1 | 0 | 2.19 | 0 |
| NXGY2 | 2021 | - | - | 0/- | 0/- | 0/- | 0/- | 0/- | 0/- | 0/- | 0/- | 0/- | 0/- | 0/- | 0/- | 0/- | 0 | 0 | 0.61 | 0 |
| SCCD2 | 2021 | - | - | 0/- | 0/- | 0/- | 0/- | 0/- | 0/- | 0/- | 0/- | 0/- | 0/- | 0/- | 0/- | 0/- | 0 | 0 | - | - |
| SDQZ | 2021 | -/0 | -/0 | 0/0 | 41.2/14.2 | 0/11.8 | 0/0 | 5.0/7.9 | 0/0 | 0/0 | 0/0 | 0/0 | 0/0 | 0/0 | 0/0 | 0/0 | 3 | 75 | 6832.26 | 50 |
| SDSG3 | 2021 | -/0 | -/50.75 | 0/0 | 0/2.5 | 3.2/6.6 | 0/0 | 0/1.7 | 0/0 | 0/0 | 0/4.3 | 0/0 | 0/0 | 0/0 | 0/0 | 0/0 | 5 | 3.23 | 1606.15 | 31.7 |
| SDSG4 | 2021 | - | - | 0/- | 61.1/- | 2.8/- | 0/- | 6.3/- | 0/- | 0/- | 1.6/- | 0/- | 0/- | 0/- | 0/- | 0/- | 4 | 84.21 | 36.84% | 76.1 |
| SDSG5 | 2021 | - | - | 0/- | 1.8/- | 1.8/- | 0/- | 31.7/- | 0/- | 0/- | 0/- | 0/- | 0/- | 0/- | 0/- | 0/- | 3 | 50 | 3103.07 | 33.9 |
| SDSG6 | 2021 | - | - | 0/- | 0/- | 1.9/- | 0/- | 88.0/- | 0/- | 0/- | 0/- | 0/- | 0/- | 0/- | 0/- | 0/- | 2 | 100 | 23612.38 | 55.4 |
| YNKM2 | 2021 | - | - | 0/- | 5.0/- | 5.0/- | 0/- | 0/- | 0/- | 0/- | 57.81/- | 0/- | 0/- | 6.25/- | 0/- | 0/- | 4 | 77.42 | 2639.36 | 47.4 |
| BJMY | 2024 | - | - | - | - | - | - | - | - | - | - | - | - | - | - | - | - | - | 5542.88 | - |
| SDSG7 | 2024 | - | - | - | - | - | - | - | - | - | - | - | - | - | - | - | - | - | 6106.12 | - |
| BJHD4 | 2024 | - | - | 0/- | 0/- | 15.0/- | 0/- | 0/- | 0/- | 0/- | 0/- | 0/- | 0/- | 0/- | 0/- | 0/- | 1 | 15.00 | 3.08 | 4.17 |
| BJDX5 | 2024 | - | - | 0/- | 0/- | 27.08/- | 0/- | 0/- | 0/- | 0/- | 6.25/- | 0/- | 6.25/- | 0/- | 0/- | 0/- | 3 | 58.33 | 2036.90 | 39.22 |
| BJDX6 | 2024 | -/0 | -/0 | 0/0 | 0/0 | 64.58/89.8 | 0/0 | 0/0.38 | 0/0 | 0/0 | 0/0 | 27.08/0 | 56.25/11.02 | 0/0 | 0/0 | 0/0 | 4 | 100.00 | 7914.83 | 77.97 |
| BJDX7 | 2024 | - | - | 0/- | 0/- | 85.71/- | 0/- | 0/- | 0/- | 0/- | 9.09/- | 0/- | 9.09/- | 0/- | 0/- | 0/- | 2 | 100.00 | 3922.24 | 64.86 |
| GXNN | 2024 | -/51.26 | -/0 | 0/0 | 0/0 | 39.58/31.02 | 0/0 | 0/2.09 | 0/0 | 0/0 | 0/0 | 0/0 | 16.7/18.06 | 0/0 | 0/0 | 0/0 | 4 | 66.67 | 1498.59 | 27.66 |
| GZGY | 2024 | - | - | 0/- | 0/- | 70.83/- | 0/- | 0/- | 0/- | 0/- | 2.08/- | 2.08/- | 47.92/- | 0/- | 0/- | 0/- | 4 | 91.67 | - | - |
| HNCS2 | 2024 | -/0 | -/0 | 0/0 | 0/0 | 35.42/23.53 | 0/0 | 0/0 | 0/0 | 0/0 | 0/0 | 0/0 | 5.88/1.91 | 0/0 | 0/0 | 0/0 | 2 | 64.71 | 30.13 | 18.87 |
| JXYC | 2024 | - | - | 0/- | 0/- | 10.42/- | 0/- | 0/- | 0/- | 0/- | 0/- | 0/- | 8.33/- | 0/- | 0/- | 0/- | 2 | 20.83 | - | - |
| LNDD | 2024 | -/0 | -/0 | 0/0 | 0/0 | 100/99.01 | 0/0 | 0/0 | 0/0 | 0/0 | 56.82/38.38 | 0/0 | 15.91/13.24 | 0/0 | 0/0 | 0/0 | 3 | 100.00 | 3364.80 | 60.94 |
| QHHD | 2024 | -/0 | -/0 | 0/0 | 2.08/0 | 75/71.16 | 0/0 | 0/0 | 0/0 | 0/0 | 34.78/33.76 | 0/0 | 0/0 | 0/0 | 0/0 | 0/0 | 3 | 100.00 | 4590.82 | 55.56 |
| SCCD3 | 2024 | - | - | 0/- | 0/- | 60.42/- | 0/- | 0/- | 0/- | 0/- | 6.52/- | 0/- | 13.04/- | 0/- | 0/- | 0/- | 3 | 82.61 | - | - |
| SDSG8 | 2024 | - | - | 27.08/- | 0/- | 0/- | 0/- | 0/- | 10.87/- | 0/- | 0/- | 0/- | 0/- | 0/- | 0/- | 0/- | 1 | 65.22 | - | - |
| SDWF | 2024 | - | - | 0/- | 2.08/- | 10.42/- | 0/- | 72.92/- | 0/- | 0/- | 2.08/- | 0/- | 0/- | 0/- | 0/- | 0/- | 4 | 100.00 | - | - |
| YNKM3 | 2024 | - | - | 0/- | 93.75/- | 0/- | 0/- | 0/- | 0/- | 0/- | 0/- | 0/- | 0/- | 0/- | 0/- | 43.75/- | 1 | 100.00 | - | - |
| YNKM4 | 2024 | - | - | 0/- | 0/- | 0/- | 0/- | 0/- | 0/- | 0/- | 100/- | 0/- | 0/- | 0/- | 0/- | 0/- | 1 | 100.00 | - | - |
| YNYX | 2024 | -/0 | -/0 | 0/0 | 66.5/60.87 | 0/1.97 | 0/0 | 0/0 | 0/0 | 0/0 | 28.26/24 | 0/0 | 2.17/0 | 0/0 | 6.52/11.56 | 10.87/0 | 3 | 100.00 | 2956.60 | 55.56 |
| ZJHZ1 | 2024 | -/18.38 | -/0 | 0/0 | 0/0 | 83.33/78.21 | 0/0 | 0/0.68 | 0/0 | 0/0 | 10.42/9.72 | 0/0 | 20.83/18.65 | 0/0 | 0/0 | 0/0 | 5 | 100.00 | 6917.85 | 73.33 |
| ZJHZ2 | 2024 | -/12.43 | -/0 | 0/0 | 0/0 | 79.17/76.89 | 0/0 | 0/0 | 0/0 | 0/0 | 0/0 | 0/0 | 2.08/2.0 | 0/0 | 0/0 | 0/0 | 3 | 95.83 | 1884.11 | 29.79 |
| LabS | 2018 | -/0 | -/0 | -/0 | -/0 | -/0 | -/0 | -/0 | -/0 | -/0 | -/0 | -/0 | -/0 | -/0 | -/0 | -/0 | 0 | - | 1.21 | 0 |
| LabR | 2018 | -/0 | -/0 | 0/0 | 0/0 | 100/100 | 0/0 | 0/0 | 0/0 | 0/0 | 100/100 | 0/0 | 0/0 | 0/0 | 0/0 | 0/0 | 2 | 100 | 54335.80 | 84 |

**Table S3 Laboratory selection and cross-resistance study of *Tetranychus* *urticae***

| Acaricide | Population | Regression (y=) | LC_50_ (95% CI) / (mg/L) | r | RR |
| --- | --- | --- | --- | --- | --- |
| Cyetpyrafen | F1 | 1.96+2.39x | 18.58 (14.85~22.23) | 0.97 | 1.0 |
|  | F32, 16 times of selection, 2019.4.2 | 1.47+1.91x | 70.94 (58.36~92.35) | 0.99 | 3.8 |
|  | F54, 28 times of selection, 2019.9.19 | 1.7252x+0.1483 | 649.12 (513.35~895.95） | 0.98 | 34.9 |
|  | F60, 30 times of selection, 2019.11.27 | 0.8799x+2.1227 | 1862.40 (1112.58~5657.60) | 0.96 | 100.2 |
|  | F62, 31 times of selection, 2019.12.25 | 2.0722x-2.6048 | 4675.77 (3477.70~10856.77） | 0.99 | 251.7 |
|  | F66, 33 times of selection, 2020.4 LabR | 2.2271 x -5.5457 | 54335.8 | 0.88 | 2924.4 |
|  | Unselected for 66 generations, LabS | 3.67 x+4.69 | 1.21 (0.29~1.89） | 0.94 | - |
| Cyenopyrafen | LabS | 2.43x+3.60 | 3.76 (3.09~4.99） | 0.94 | 1.0 |
|  | LabR | 1.05x-0.16 | 79917.51 (32597.48~7279096.99) | 0.78 | 21227.0 |
| Cyflumetofen | LabS | 3.25 x+2.36 | 6.47 (3.58~8.86) | 0.95 | 1.0 |
|  | LabR | 0.62x+1.52 | 452003.19 | 0.97 | 69902.4 |
| *Pyridaben* | LabS | 1.47x-0.25 | 3763.36 (3013.428~5446.99） | 0.97 | 1.0 |
|  | LabR | 1.01x-0.94 | 10351.97 (5899.42~71346.098） | 0.94 | 3.2 |
| Bifenazate | LabS | 2.34x+2.67 | 9.87 (8.3894~11.4358） | 0.99 | 1.0 |
|  | LabR | 2.20x+2.28 | 17.39(14.62~21.35） | 0.98 | 1.8 |

**Table S4 Sequencing information and genetic diversity of 22 populations of the two-spotted spider mite *Tetranychus* *urticae*.** N, number of individuals for pooled sequencing. π, nucleotide diversity.

| Code | Date | Resistance status | N | Raw_reads | Mapped_reads | Mean_depth | No. of SNP | π | Tajima’ D |
| --- | --- | --- | --- | --- | --- | --- | --- | --- | --- |
| SXYQ | 2017.03.14 | Unkonwn | 300 | 226,518,938 | 165,721,999 | 277.66X | 3,408,816 | 0.00638±0.00010 | 0.4210±0.0219 |
| AHHN | 2017.03.16 | Unkonwn | 50 | 174,220,938 | 168,955,686 | 208.99X | 2,619,282 | 0.00644±0.00010 | 0.3001±0.0228 |
| SDRZ | 2017.03.22 | Unkonwn | 300 | 223,695,602 | 138,326,975 | 230.89X | 2,870,693 | 0.00761±0.00011 | 0.2411±0.0150 |
| SCCD1 | 2017.03.30 | Unkonwn | 200 | 199,016,876 | 166,311,650 | 278.69X | 4,628,586 | 0.00811±0.00010 | -0.0305±0.0158 |
| HNHK | 2017.03.30 | Unkonwn | 300 | 209,078,536 | 133,365,977 | 223.88X | 2,196,442 | 0.00523±0.00010 | 0.2197±0.0232 |
| SHPD | 2017.04.13 | Unkonwn | 300 | 254,783,024 | 176,667,923 | 296.06X | 3,073,153 | 0.00650±0.00010 | 0.3568±0.0198 |
| HNCS1 | 2017.04.02 | Unkonwn | 300 | 209,671,400 | 151,838,290 | 254.52X | 4,149,620 | 0.00726±0.00010 | 0.1330±0.0190 |
| JXNC | 2017.05.08 | Unkonwn | 300 | 145,020,244 | 99,536,930 | 167.41X | 2,225,209 | 0.00552±0.00010 | 0.1681±0.0218 |
| NMHH2 | 2021.06.13 | Susceptible | 300 | 220,519,748 | 149,102,857 | 249.51X | 2,835,462 | 0.00758±0.00011 | 0.1967±0.0151 |
| BJCP4 | 2021.04.27 | Resistant | 230 | 223,631,126 | 139,731,804 | 234.65X | 2,898,958 | 0.00745±0.00011 | 0.1201±0.0155 |
| SDQZ | 2021.06.13 | Resistant | 200 | 188,858,914 | 119,965,047 | 197.17X | 3,231,043 | 0.00743±0.00011 | 0.2025±0.0180 |
| SDSG3 | 2021.04.18 | Resistant | 250 | 188,571,660 | 125,730,235 | 209.41X | 3,109,064 | 0.00687±0.00011 | 0.1560±0.0202 |
| HNCS2 | 2024.01.29 | Susceptible | 300 | 222,928,412 | 151,838,290 | 254.52X | 3,046,516 | 0.00741±0.00010 | 0.1015±0.0151 |
| BJDX6 | 2024.03.15 | Resistant | 300 | 283,068,416 | 191,429,676 | 322.60X | 3,056,055 | 0.00731±0.00010 | 0.0375±0.0153 |
| GXNN | 2024.03.31 | Resistant | 100 | 342,901,124 | 157,079,428 | 264.42X | 3,458,964 | 0.00793±0.00011 | 0.0967±0.0156 |
| LNDD | 2024.03.10 | Resistant | 250 | 306,161,484 | 176,939,299 | 296.53X | 2,723,643 | 0.00722±0.00010 | 0.2689±0.0161 |
| QHHD | 2024.02.29 | Resistant | 50 | 240,992,080 | 135,847,928 | 228.76X | 2,961,593 | 0.00767±0.00011 | 0.1533±0.0168 |
| YNYX | 2024.03.05 | Resistant | 150 | 233,843,470 | 140,711,946 | 236.87X | 2,510,601 | 0.00665±0.00010 | 0.3204±0.0192 |
| ZJHZ1 | 2024.03.15 | Resistant | 400 | 257,063,980 | 146,899,909 | 247.31X | 3,105,779 | 0.00737±0.00010 | 0.0960±0.0154 |
| ZJHZ2 | 2024.03.15 | Resistant | 400 | 210,338,514 | 132,308,752 | 222.79X | 3,306,753 | 0.00747±0.00010 | 0.0585±0.0156 |
| LabS | 2018.04 | Susceptible | 200 | 136,765,686 | 99,182,723 | 165.93X | 2,868,547 | 0.00701±0.00010 | 0.2366±0.0175 |
| LabR | 2020.05 | Resistant | 200 | 130,487,654 | 96,193,157 | 160.86X | 2,958,783 | 0.00680±0.00010 | 0.1220±0.0180 |
| Total number |  |  | 5680 |  |  |  |  |  |  |

**Table S5 Distribution of haplotypes carrying amino acid mutations in populations.** Population codes in red indicate the resistant population. Haplotype codes in blue indicates the putative ancestral haplotype, while those in red indicate those carrying the potential resistant mutations.

| Population | *sdhB* | | | | | | | | | | |  | *sdhD* | | | | | | | | | | | | | | | | | | | | | | | |
| --- | --- | --- | --- | --- | --- | --- | --- | --- | --- | --- | --- | --- | --- | --- | --- | --- | --- | --- | --- | --- | --- | --- | --- | --- | --- | --- | --- | --- | --- | --- | --- | --- | --- | --- | --- | --- |
|  | H1 | H2 | H3 | H4 | **H5** | **H6** | H7 | **H8** | H9 | H10 | **H11** |  | H1 | H2 | H3 | H4 | **H5** | **H6** | **H7** | **H8** | H9 | **H10** | H11 | H12 | **H13** | H14 | H15 | **H16** | **H17** | H18 | H19 | **H20** | **H21** | **H22** | H23 | H24 |
| BJCP1 | 13 | 3 | 4 | 2 |  |  |  |  |  |  |  |  | 5 | 6 | 1 | 1 |  |  |  |  |  |  |  |  |  |  |  |  |  |  |  |  |  |  |  |  |
| BJPG1 | 3 |  |  | 9 |  |  |  |  |  |  |  |  | 1 | 1 | 1 |  |  |  |  |  |  |  | 3 | 1 |  |  |  |  |  |  |  |  |  |  |  |  |
| BJCP2 | 7 |  | 4 | 16 |  |  |  |  |  |  |  |  |  | 3 | 13 |  |  |  |  |  |  |  |  |  |  |  |  |  |  |  |  |  |  |  |  |  |
| BJHD1 | 11 |  | 1 | 4 |  |  |  |  |  |  |  |  |  | 19 |  |  |  |  |  |  |  |  |  |  |  |  |  |  |  |  |  |  |  |  |  |  |
| BJPG2 | 10 |  |  | 9 |  |  | 8 |  |  |  |  |  | 6 | 10 | 1 |  |  |  |  |  |  |  | 3 |  |  |  |  |  |  |  |  |  |  |  |  |  |
| BJTZ1 | 16 |  | 3 | 8 |  |  |  |  |  |  |  |  | 12 | 16 | 1 |  |  |  |  |  |  |  | 5 |  |  |  |  |  |  |  |  |  |  |  |  |  |
| HNCS1 |  |  | 1 | 26 |  |  |  |  |  |  |  |  | 4 | 7 |  |  |  |  |  |  |  |  | 15 |  |  | 3 |  |  |  |  |  |  |  |  |  |  |
| HNHK | 4 |  |  | 16 |  |  |  |  |  |  |  |  | 4 | 14 |  |  |  |  |  |  |  |  | 4 | 1 |  |  |  |  |  |  |  |  |  |  |  |  |
| SCCD1 | 10 |  | 1 | 8 |  |  | 1 |  |  |  |  |  | 7 | 1 | 4 |  |  |  |  |  |  |  | 1 |  |  |  |  |  |  |  |  |  |  |  |  |  |
| SDRZ | 10 |  | 2 | 11 |  |  | 3 |  |  |  |  |  | 2 | 13 | 1 |  |  |  |  |  |  |  | 2 |  |  |  |  |  |  |  |  |  |  |  |  |  |
| SDSG1 | 3 |  |  | 1 |  |  |  |  |  |  |  |  | 7 |  | 1 |  |  |  |  |  |  |  |  |  |  |  |  |  |  |  |  |  |  |  |  |  |
| SHPD | 1 |  |  | 22 |  |  |  |  |  |  |  |  |  | 5 | 18 |  |  |  |  |  |  |  |  |  |  |  |  |  |  |  |  |  |  |  |  |  |
| BJHD2 | 6 |  |  | 1 |  |  |  |  |  |  |  |  | 1 |  |  |  |  |  |  | **11** |  |  |  |  |  |  |  |  |  |  |  |  |  |  |  |  |
| HNZZ | 7 |  | 4 | 10 |  |  |  |  |  |  |  |  | 2 | 5 | 8 |  |  |  |  |  |  |  | 2 |  |  |  | 1 |  |  |  |  |  |  |  |  |  |
| LNSY | 3 |  |  | 6 |  |  | 1 |  |  |  |  |  | 3 | 6 |  |  |  |  |  |  |  |  |  |  | **1** |  |  |  | **3** |  |  |  |  |  |  |  |
| SDQD | 2 | 10 | 1 | 4 |  |  | 5 |  |  |  |  |  | 1 | 1 | 2 |  |  |  |  |  |  |  | 14 |  |  |  |  |  |  | 1 |  |  |  |  |  |  |
| SDSG2 | 12 |  |  | 4 |  |  |  |  |  |  |  |  | 2 |  |  |  |  |  |  |  |  |  |  |  |  |  |  |  |  |  |  | **10** |  |  |  |  |
| SXAK | 19 |  |  | 2 |  |  |  |  |  |  |  |  |  | 20 |  |  |  |  |  |  |  |  | 2 |  |  |  |  |  |  |  |  |  |  |  |  |  |
| YNKM1 | 6 | 1 | 1 | 12 |  |  |  |  |  |  |  |  | 1 | 2 | 7 |  |  |  |  |  |  |  | 6 |  |  |  |  |  |  |  |  |  |  |  |  |  |
| **BJCP4** | 2 |  | 1 | 5 | **6** |  |  |  |  |  |  |  |  | 1 | 1 |  | **20** |  |  |  |  |  |  |  |  |  |  |  |  |  |  |  |  |  |  |  |
| BJHD3 | 2 |  |  | 11 |  | **2** |  |  |  |  |  |  |  |  | 4 |  |  |  |  | **15** |  |  |  |  |  |  |  |  |  |  |  |  |  |  |  |  |
| BJPG3 |  |  |  |  | **2** | **11** |  |  |  |  |  |  | 3 | 8 | 2 |  |  |  |  |  |  | **1** | 2 |  | **4** |  |  |  |  |  |  |  |  |  |  |  |
| BJTZ2 | 1 |  |  |  | **2** | **3** |  |  |  |  |  |  | 5 | 7 | 4 |  |  |  |  |  |  | **1** |  |  |  |  |  |  |  |  |  |  |  |  |  |  |
| NMHH2 | 14 |  | 1 | 1 |  |  |  |  |  |  |  |  |  |  |  |  |  |  |  |  |  |  | 1 |  |  |  |  |  |  |  |  |  |  |  |  |  |
| NXGY2 | 20 |  | 1 | 2 |  |  | 2 |  |  |  |  |  |  | 2 | 7 |  |  | **6** |  |  |  |  |  |  |  |  |  |  |  |  |  |  |  |  |  |  |
| SCCD2 | 4 |  | 2 | 9 |  |  | 15 |  |  |  |  |  |  | 16 |  |  |  |  |  |  |  |  |  |  |  |  |  |  |  |  |  |  |  |  |  |  |
| **SDQZ** | 4 |  | 1 |  |  |  |  | **2** |  |  |  |  | 8 | 2 | 2 |  |  |  |  |  |  |  |  |  |  |  |  |  |  |  | 4 |  |  |  |  |  |
| **SDSG3** | 9 |  | 1 | 20 |  | **1** |  |  |  |  |  |  | 24 |  | 3 |  |  |  |  |  |  |  |  |  |  |  |  |  |  |  |  |  |  |  |  |  |
| SDSG4 |  |  |  | 2 |  |  |  | **7** |  |  |  |  | 7 | 2 | 3 |  |  |  |  |  |  |  | 3 |  |  | 3 |  |  |  |  |  |  | **1** |  |  |  |
| SDSG5 | 20 |  | 1 | 5 |  |  |  |  |  |  |  |  | 8 |  | 1 |  |  |  |  |  |  |  |  |  |  |  |  |  |  |  |  | **6** |  |  |  |  |
| SDSG6 | 3 |  | 1 | 19 |  |  |  |  | 1 |  |  |  |  |  |  |  |  |  |  |  |  |  |  |  |  |  |  |  |  |  |  |  | **14** |  |  |  |
| YNKM2 | 8 |  | 5 | 10 |  |  |  |  |  |  |  |  |  |  |  |  | **16** |  |  |  |  |  |  |  |  |  |  |  |  |  |  |  |  | **1** |  |  |
| BJDX5 | 5 |  |  | 1 |  | **1** |  |  |  |  |  |  | 1 | 4 |  |  |  |  |  |  |  |  |  |  |  |  |  |  |  |  |  |  |  |  |  |  |
| **BJDX6** |  |  |  | 2 |  | **9** |  |  |  |  |  |  |  |  |  |  |  | **1** | **1** |  |  |  |  |  |  |  |  |  |  |  |  |  |  |  |  |  |
| BJDX7 |  |  |  | 1 |  | **13** |  |  |  |  |  |  | 1 | 1 |  |  |  |  |  |  |  |  |  |  |  |  |  |  |  |  |  |  |  |  |  |  |
| BJHD4 | 3 |  |  |  |  | **1** |  |  |  |  |  |  |  |  | 6 |  |  |  |  |  | 1 |  |  |  |  |  |  |  |  |  |  |  |  |  |  |  |
| **GXNN** | 2 |  |  | 7 | **2** | **4** |  |  |  |  |  |  | 1 |  |  |  |  | **1** |  |  |  |  |  |  |  |  |  |  |  |  |  |  |  |  |  |  |
| GZGY | 2 |  |  |  | **1** | **6** |  |  |  |  |  |  | 1 |  |  |  |  |  |  |  |  |  |  |  |  |  |  |  |  |  |  |  |  |  |  |  |
| **HNCS2** |  |  |  | 6 |  | **2** |  |  |  |  |  |  | 1 | 4 | 1 |  |  |  |  |  |  |  |  |  |  |  |  |  |  |  |  |  |  |  |  |  |
| JXYC | 7 |  |  | 3 |  | **2** | 2 |  |  |  |  |  | 5 | 5 |  |  |  |  |  |  |  |  |  |  |  |  |  | **1** |  |  |  |  |  |  |  |  |
| **LNDD** |  |  |  |  | **10** | **3** |  |  |  |  |  |  |  | 1 |  |  |  |  |  |  |  |  |  |  |  |  |  |  | **3** |  |  |  |  |  |  |  |
| **QHHD** |  | 1 |  |  |  | **12** |  |  |  |  |  |  |  | 1 | 2 |  |  |  |  |  |  |  |  |  |  |  |  |  | **2** |  |  |  |  |  |  |  |
| SCCD3 | 1 |  |  | 2 |  | **10** |  |  |  |  |  |  | 1 | 2 | 1 |  |  |  |  |  |  |  | 1 |  |  |  |  |  |  |  |  |  |  |  |  |  |
| SDSG8 | 12 |  |  |  |  |  |  |  |  | 2 |  |  |  | 5 | 3 |  |  |  |  |  |  |  |  |  |  |  |  |  |  |  |  |  |  |  |  |  |
| SDWF | 9 |  |  |  |  |  |  |  |  |  |  |  |  |  |  |  |  |  |  |  |  |  |  |  |  |  |  |  |  |  |  | **10** |  |  |  |  |
| YNKM3 |  |  |  | 4 |  |  |  | **5** |  |  | **5** |  |  | 1 |  |  | **23** |  |  |  |  |  |  |  |  |  |  |  |  |  |  |  |  |  | 4 |  |
| YNKM4 | 9 |  |  |  |  |  |  |  |  |  |  |  |  |  |  |  |  |  |  |  |  |  |  |  |  |  |  |  |  |  |  |  |  |  |  |  |
| **YNYX** |  |  |  | 2 |  |  |  | **2** |  |  | **1** |  | 6 |  |  |  | **1** |  |  |  |  |  |  |  |  |  |  |  |  |  |  |  |  |  |  |  |
| **ZJHZ1** |  |  |  |  |  | **18** |  |  |  |  |  |  |  |  |  |  | **1** | **1** |  |  |  |  |  |  |  |  |  |  |  |  |  |  |  |  |  |  |
| **ZJHZ2** |  |  |  |  |  | **11** |  |  |  |  |  |  |  | 5 | 1 |  |  |  |  |  |  |  | 3 |  |  |  |  |  |  |  |  |  |  |  |  |  |
| ZJXS1 | 14 |  | 1 | 17 |  |  | 4 |  |  |  |  |  | 2 | 11 | 4 | 1 |  |  |  |  |  |  | 10 |  |  |  |  |  |  |  |  |  |  |  |  | 1 |
| **LabR** |  |  |  |  |  | **24** |  |  |  |  |  |  |  |  |  |  |  |  |  |  |  | **32** |  |  |  |  |  |  |  |  |  |  |  |  |  |  |

**Table S6 Correlation between eight mutations on SDH genes and level of resistance (survival percentages under 1000 mg/L)**

| **Mutations** | **Regression (y =)** | **Coefficient (R2)** |
| --- | --- | --- |
| sdhB_I260T | 0.526x + 0.358 | 0.0927 |
| sdhB_I260V | -0.00222x + 0.4 | 0.0031 |
| sdhD_R119C | 0.496x + 0.318 | 0.198 |
| sdhD_R119L | 0.808x + 0.353 | 0.0981 |
| sdhD_R119G | 1.47x + 0.381 | 0.0647 |
| sdhD_R119H | 0.769x + 0.394 | 0.00107 |
| sdhD_R119P | -0.474x + 0.413 | 0.0669 |
| sdhD_D116G | 0.4967x + 0.2937 | 0.1605 |
| Predominant resistant allele | 0.757x + 0.0401 | 0.703 |
| Individuals with at least one resistant allele | 0.649x - 0.0088 | 0.747 |
| Individuals with at least one homozygous resistant genotype | 0.702x + 0.132 | 0.606 |

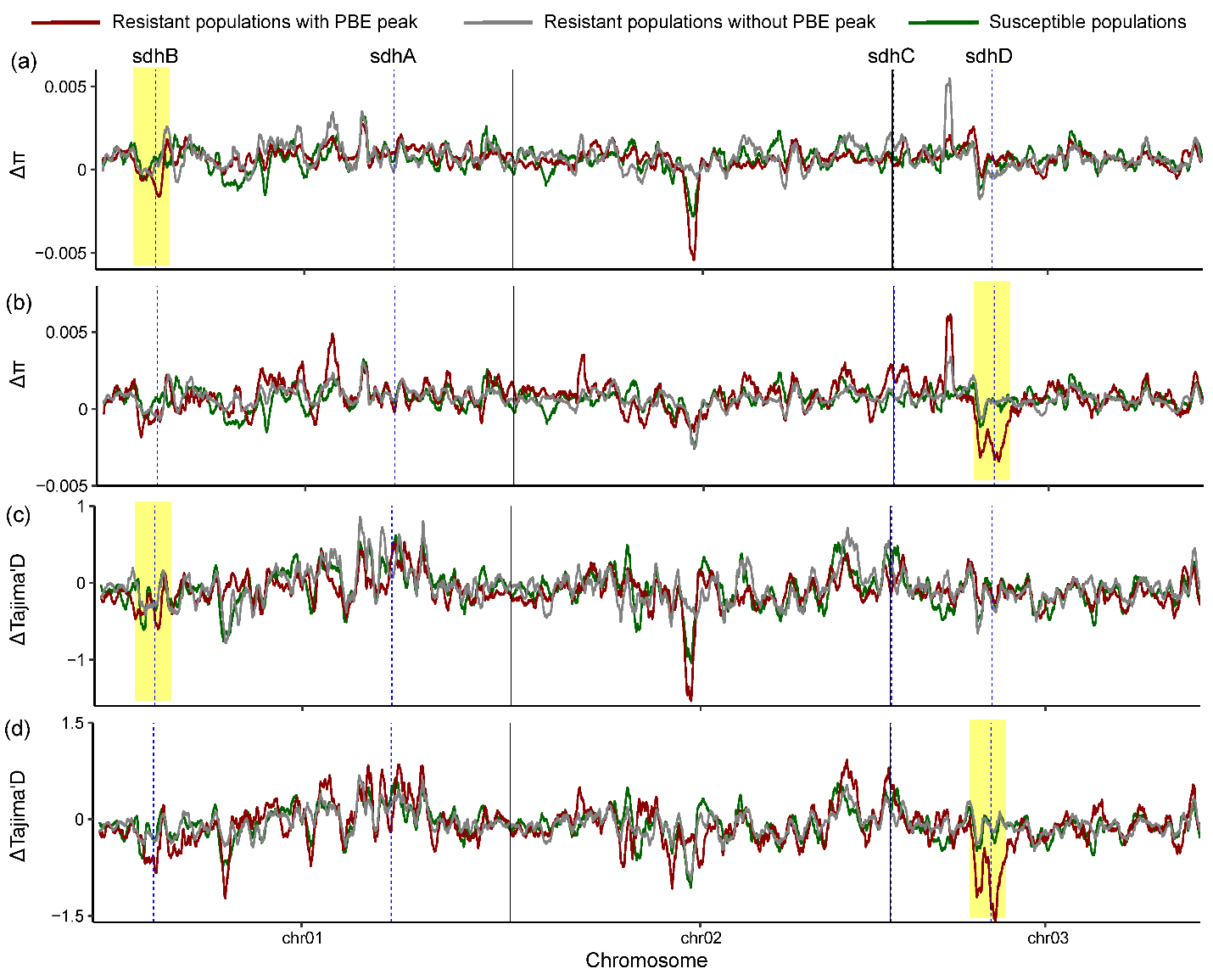

**Fig. S1 Moving average of delta nucleotide diversity (a - b) and delta Tajima’D values (c - d) along chromosomes.** Delta π and Tajima's D represent the difference in nucleotide diversity (π) and Tajima's D value, respectively, between three field population groups collected before the commercial release of cyetpyrafen and the field populations collected after the release. Lines with different colors represent moving average of 100 genomic windows of 5 kbp wide in three population groups. The red lines indicate four (a, c) or one field-collected resistant population (b, d) with a clear peak around sdhB or sdhD genes in Fig. 2b, while the grey points and lines indicate seven or ten field-collected resistant populations without a peak around sdhB or sdhD genes in Fig. 2b. The green lines represent two susceptible populations. The vertical dashed lines show the position of SDH genes. Signals of selection sweep, characterized by decreased nucleotide diversity and Tajima's D values, were observed in resistant populations (red lines), with the selection signal identified by PBE analysis in Fig. 2b.

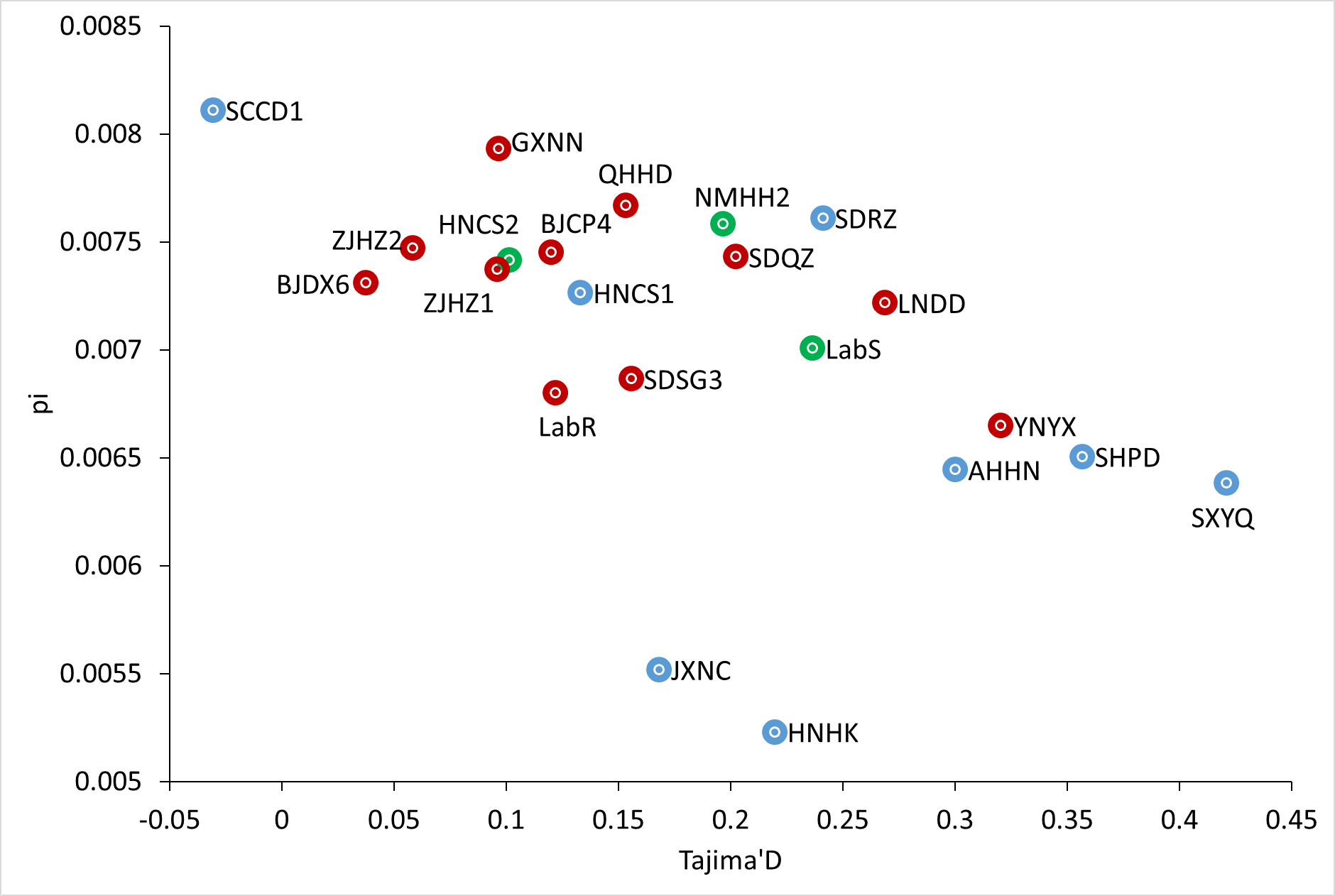

**Fig. S2 Scatter plot of average genome-wide nucleotide diversity (π) and Tajima's D per population.** The red points, green points, and blue points indicate resistant populations, susceptible populations, and populations collected before the commercial release of cyetpyrafen, respectively.

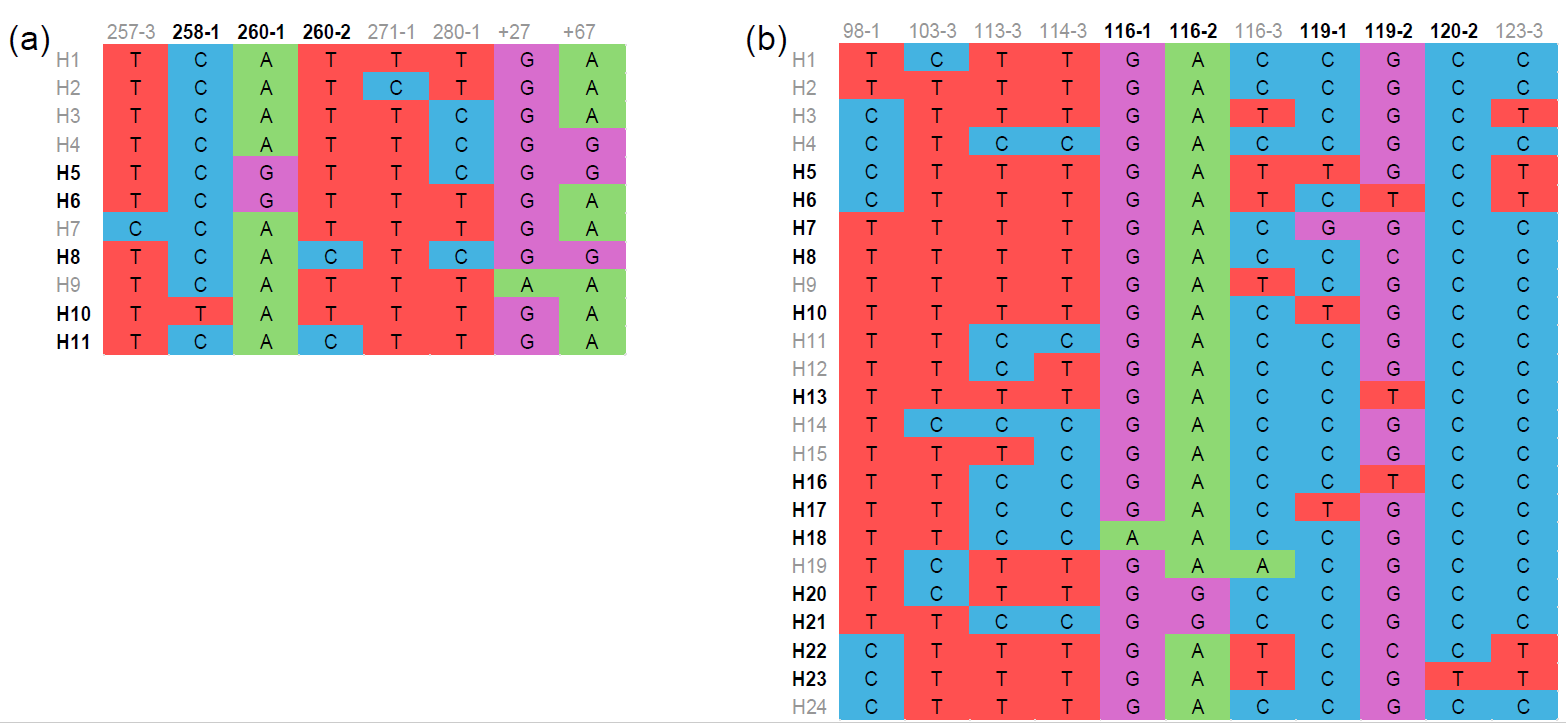

**Fig. S3 Haplotypes of sdhB (a) and sdhD (b) genes.** (a) and (b), alignment of haplotypes for sdhB and sdhD genes. Only variant sites are displayed. Nucleotide sites are numbered according to the amino acid position they coded for, followed by the number of the codon positions. The codes of haplotypes carrying nonsynonymous mutation sites are in dark, while codes of haplotypes carrying synonymous mutation sites are in grey.

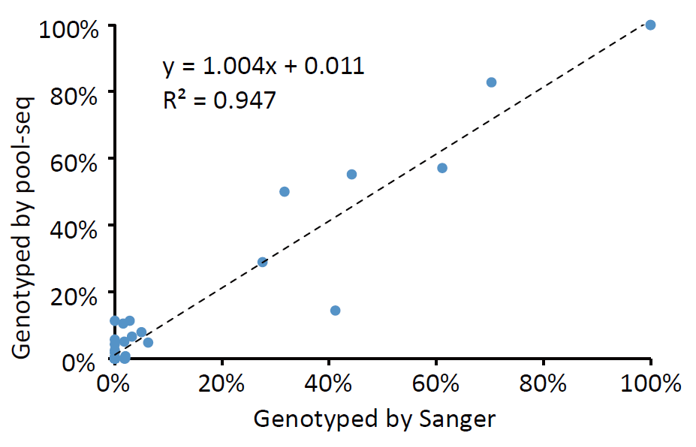

**Fig. S4** Correlation between allele frequencies genotyped by Sanger sequencing and pool-seq.
